# NERVE 2.0: boosting the New Enhanced Reverse Vaccinology Environment via artificial intelligence and a user-friendly web interface

**DOI:** 10.1101/2024.11.13.623451

**Authors:** Andrea Conte, Nicola Gulmini, Francesco Costa, Matteo Cartura, Felix Bröhl, Francesco Patanè, Francesco Filippini

## Abstract

**Background:** Vaccines development in this millennium started by the milestone work on *Neisseria meningitidis B*, reporting the invention of Reverse Vaccinology (RV), which allows to identify vaccine candidates (VCs) by screening bacterial pathogens genome or proteome through computational analyses. When NERVE (New Enhanced RV Environment), the first RV software integrating tools to perform the selection of VCs, was released, it prompted further development in the field. However, the problem-solving potential of most, if not all, RV programs is still largely unexploited by experimental vaccinologists that impaired by somehow difficult interfaces, requiring bioinformatic skills.

**Results:** We report here on the development and release of NERVE 2.0 (available at: https://nerve-bio.org) which keeps the original integrative and modular approach of NERVE, while showing higher predictive performance than its previous version and other web-RV programs (Vaxign and Vaxijen). We renewed some of its modules and added innovative ones, such as *Loop Razor*, to recover fragments of promising vaccine candidates or *epitopepredict* for the epitope prediction binding affinities and population coverage. Along with two newly built AI (Artificial Intelligence)-based models: ESPAAN and Virulent. To improve user-friendliness, NERVE was shifted to a tutored, web-based interface, with a noSQL-database to consent the user to submit, obtain and retrieve analysis results at any moment.

**Conclusions:** With its redesigned and updated environment, NERVE 2.0 allows customisable and refinable bacterial protein vaccine analyses to all different kinds of users.

## 1. Background

In the early 2000s, the growing availability of genomic data and the development of more performing bioinformatics tools led to a revolution in vaccinology, i.e., to the birth of *reverse vaccinology* (RV) [1]. Starting with the work on *Neisseria meningitidis group B* by Rino Rappuoli’s team [2], RV has enhanced the capacity to identify VCs by replacing several experimental tasks. This is possible via in-silico prediction steps on the genome and/or proteome of the pathogen of interest, with consequent time and cost benefits. Afterwards, RV has been applied to other pathogenic bacterial species. However, the first bioinformatics-driven approaches were pathogen-tailored and poorly generalizable [3]. To tackle the need to standardise VCs search process, the first published open-source RV platform was NERVE (*New Enhanced Reverse Vaccinology Environment)* [4], only for Linux users. It is based on a modular structure, and it extracts relevant information from a given pathogen proteome through six analytical steps, comprehending different bioinformatic tools (see Additional figure 1). These data, saved in a MySQL table, are then used to infer the presence of protein vaccine candidates (PVCs), which are collected in an HTML table.

**Figure 1.**
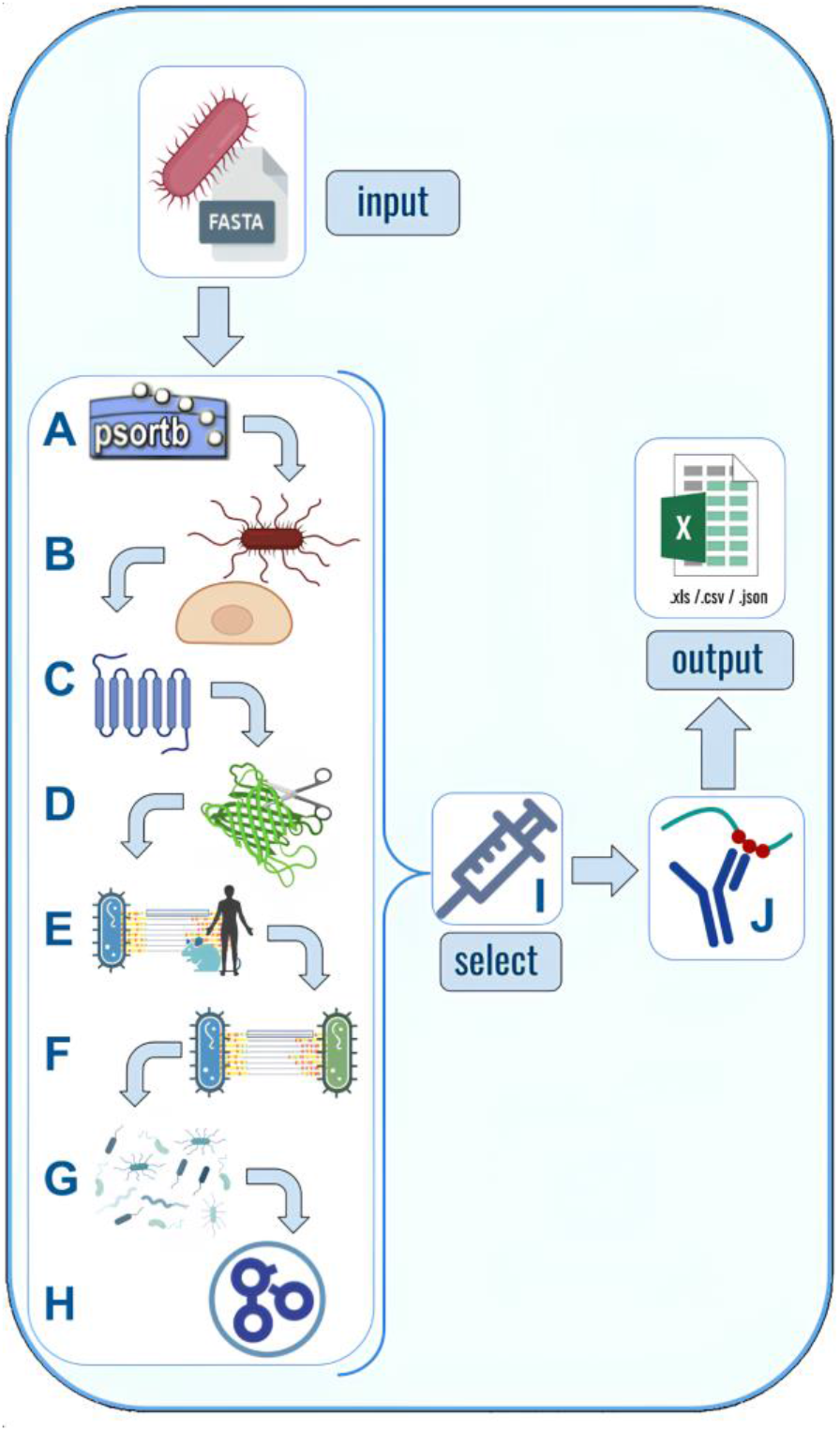
NERVE 2.0 working structure. Bacterial protein sequences are provided as an input FASTA proteome and undergo eight analytical steps: (A) *Subcelloc* predicts protein subcellular localization, (B) *Adhesin* returns the probability of a protein to be an adhesin, (C) *Tmhelices* predicts protein topology, (D) *Loop-Razor* rescues membrane proteins reduced to their extracellular fragments, (E) *Autoimmunity and Mouse Immunity* which find respectively matches between the pathogen under analysis and human or mice proteomes (F) *Conservation* which detects conserved proteins between two input bacterial strains, (G) *Virulent* to infer presence of virulence factors and (H) *Annotation to predict protein function*. Then, the *Select* module (I) filters out PVCs, which meet specific requirements. Output results can be downloaded in .json, .csv, or .xlsx format. *Epitope* prediction (J) is performed after the Select module. Created with BioRender.com.

Inspired by NERVE, further RV applications have been developed, including new and updated tools to improve VCs identification [3]. According to their working structure, RV programs can be classified into these two categories [9]: filter-based and machine learning (ML)-based. In the first ones, protein sequences are analysed by different tools to obtain useful information, such as their probability of being an adhesin or their subcellular localization. Then, this is passed to a decision tree that uses a priori cut-offs to select PVCs. NERVE [4], Jenner-predict [10], VacSol [11] and Vaxign [12] fall into this category.

In the second category, some specific protein sequence features are extracted and directly fed to a ML model. These features may include physicochemical descriptors, such as amino acid composition or hydrophobicity propensity. Next, an artificial neural network (ANN) is often adopted to classify input proteins in PVCs and not-PVCs, solving a common binary classification task. Examples of these programs are VaxiJen [13] and Bowman-Heinson [14,15].

Differently from the cited ones, ReVac [16] uses a scoring system that ranks all PVCs from the most to the least likely one. It accepts bacterial proteomes and genomes as inputs, which are analysed by several bio-tools integrated in the Ergatis platform [17] and grouped according to the analysed feature, enabling parallel computations. This redundancy, as the authors specify, leads to more confident predictions. Nevertheless, the complex structure of ReVac represents a drawback because it requires unclear software dependency installation and scarce documentation, representing a major disadvantage for most of the users.

Despite recent and remarkable improvements in RV applications, such as the evaluation of virulence factors as PVCs [11] or population coverage predictions of pathogen input proteins [12], one of their main limitations remains the difficulty of installation and use. Most of the cited programs are not readily accessible because their installation procedures are often challenging, due to unclear instructions or little support by the developers. Instead, accessibility to a broad plethora of users should be an essential goal of RV applications [9]. NERVE 1.0 installation was rather arduous as well because of the multiple dependencies which required manual downloading and configuration, and it is no longer supported as most of its Perl libraries are now obsolete.

We upgraded NERVE to tackle the usability and accessibility pitfalls common to many RV programs. We renewed most modules (also named components) and included new AI (Artificial Intelligence) ones. In the realm of vaccine development, advanced computational pipelines are now utilising AI systems, which provide great improvements in terms of accuracy and speed allowing also a better comprehensive analysis of entire proteomes, compared to non-AI methods [41].

## 2. Implementation

NERVE 1.0 was meant to be an environment, gathering different tools and making them collaborate to obtain a final PVCs ranking. The same approach is maintained in version 2.0.

The first major difference, compared to the original version, is the programming language: Perl was replaced by Python, especially for libraries availability, e.g. TensorFlow [18] for the modules using deep learning models. Regarding the input, FASTA files can be automatically retrieved from UniProt [19], in addition to manual uploading. They are also subject to a *quality check* to exclude sequences presenting anomalous characters or non-conventional amino acids.

The results are saved in a .csv table, which is easier to visualise and download. They are also permanently stored on the server-side and can be accessed at any moment by the user. The output is then available in three formats: .csv, .xls, and .json.

Part of NERVE 2.0 modules were substituted with new packages created ad hoc. Python-TMHMM, a protein topology predictor [20] was adapted to fit the new NERVE pipeline. The adhesin predictor SPAAN [6] was completely renewed adopting a different ML architecture and using a new training dataset. The same model was also trained on different datasets to obtain *Virulent*, a module for the prediction of virulent factors. In addition, ML was considered for protein function prediction, performed by DeepFri [34]. Four new components have been added to NERVE 2.0, namely *LoopRazor, Virulent, Mouse Immunity* and *Epitope prediction*. A brief description of each component is provided hereafter, while Figure 1 summarises NERVE

### 2.0 overall structure

Users interacting with the web interface or using NERVE 2.0 stand-alone version (see *Availability of data and materials*), can decide to optionally activate only some of these components and modify specific parameters allowing customisable PVCs search.

### 2.1. Subcelloc module

*Subcelloc* predicts input protein subcellular localizations and aims at finding surface-exposed proteins, which are ideal vaccine targets [6]. For this purpose, this module uses PSORTb 3.0 [21], which shows improved precision and recall with respect to the version 2.0, required for the original NERVE. PSORTb 3.0 predicts the subcellular localization given the bacterial Gram type, which is a classification based on bacterial membrane staining [21]. Predictions equal or above the threshold of 7.5 are considered valid in accordance with the PSORTb 3.0 documentation [21].

### 2.2. ESPAAN, adhesin prediction module

Bacterial adhesins are surface-associated virulence factors that play a crucial role during the first steps of infection, mediating attachment to host cells [22]. Because they are surface bound and required for the infection to occur, they can be readily targeted by the immune system thus representing valid PVCs [22]. A list of known adhesive domains derived from the literature [23] was used to construct a dataset via jackhammer search on reference eubacteria proteomes using the HMMER web server (HMMER 3.3.2) [24]. A set of putative non-adhesin proteins was derived from a search on the SwissProt section (manually reviewed and curated proteins) of Uniprot [19] considering bacterial proteins with non-adhesin related keywords. Redundant sequences (25% identity threshold) were removed from both datasets with CD-hit [25] obtaining 2700 adhesin proteins. A subset of non-adhesin proteins was randomly selected to match the size of the adhesin dataset. Proteins with local similarity above 25% with proteins used to tune the *Select* module (see section 2.12.), as measured with BLASTp [8], have also been removed.

A neural network based on a 10-unit Dense layer was implemented for adhesins identification. The network takes some protein features as input, and it is trained to correctly classify if a protein is an adhesin or not. Cross-entropy loss is used. The features considered are computed with the Python package iFeature [26] and consist of amino acid composition (AAC), dipeptide composition (DPC), composition (CTDC), transition (CTDT), and distribution (CTDD), as similarly done in PathoFact, a tool for the prediction of virulence factors and antimicrobial genes [27]. Since every sequence has a (20+400+39+39+195=693)-dimensional feature vector, we performed Principal Component Analysis (PCA) to reduce the dimensionality. We found 400 features to be sufficient to explain the variation observed in the dataset, which was split into 50% training, 25% validation and 25% test set. The training was performed for 120 epochs. To test ESPAAN performance, the main evaluation metrics (shown in Figure 2 and Table 1) were calculated on its test set.

**Table 1.** Evaluation metrics of ESPAAN and SPAAN, calculated on ESPAAN test set using a dedicated python script with TensorFlow. The test set consists of 692 positive and 658 negative sequences. 5 sequences from the positive set and 25 from the negative sets were filtered out by SPAAN and therefore not considered to calculate its metrics.

| Metric | ESPAAN | SPAAN |
| --- | --- | --- |
| Accuracy | 0.932 | 0.669 |
| Sensitivity (Recall) | 0.916 | 0.566 |
| Precision | 0.949 | 0.737 |
| F1-score | 0.932 | 0.640 |
| False Positive Rate (FPR) | 0.052 | 0.220 |
| False Negative Rate (FNR) | 0.084 | 0.434 |
| True Negative Rate (TNR) | 0.948 | 0.780 |
| False Discovery Rate (FDR) | 0.051 | 0.263 |
| Negative Predictive Value (NPV) | 0.915 | 0.624 |

**Figure 2.**
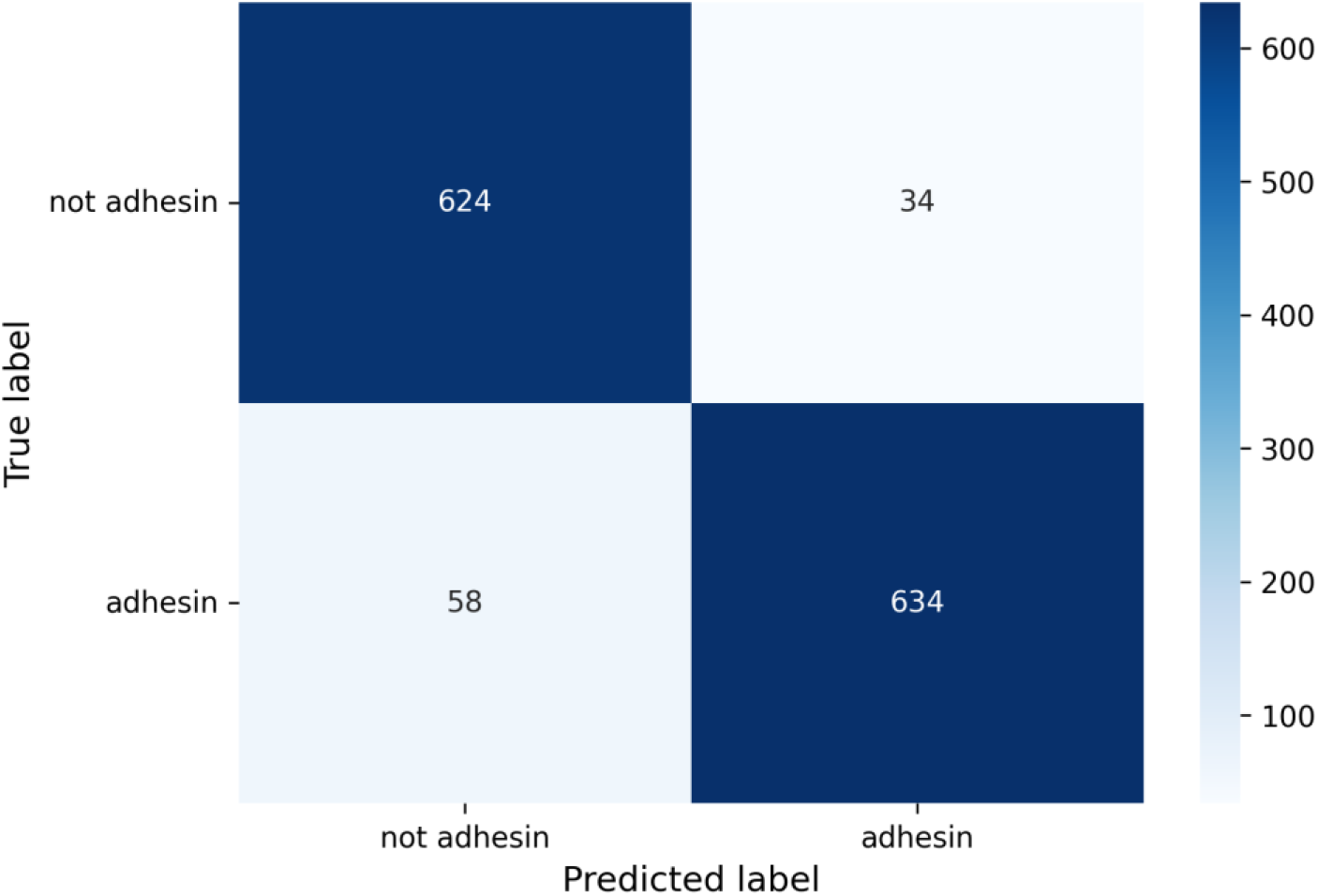
ESPAAN confusion matrix. A 2×2 matrix has been considered for this binary classification problem (adhesin / non-adhesin), setting PAD (probability of being adhesin) = 0.5 as threshold.

ESPAAN shows very good performances. Precision and recall both have high values (> 0.9), and therefore F1-score is high, too (0.932). Additionally, probabilities of finding false positives and negatives are quite low (0.052 and 0.084 respectively). The 0.932 recorded accuracy demonstrates that ESPAAN makes overall correct predictions with few exceptions. In addition, ESPAAN was also benchmarked against its predecessor, SPAAN^1^, demonstrating a notable superiority (Table 1).

To better assess ESPAAN performances and to avoid overfitting, a 3-fold cross validation was performed with TensorFlow. As evidenced by data presented in Table 2, ESPAAN also shows very high mean validation metrics (> 0.9) with very low related standard deviations. This reconfirms the results of the previous demonstration.

**Table 2.** 3-fold cross validation of ESPAAN with related obtained metrics.

| Metric | Value | Std deviation |
| --- | --- | --- |
| Mean validation accuracy | 0.929 | 0.002 |
| Mean validation recall | 0.932 | 0.005 |
| Mean validation precision | 0.914 | 0.002 |

**S**See Additional file 1 in *Supplementary Material* and *Availability of data and materials* for an overall comprehension of ESPAAN tests data.

### 2.3. TMhelices module

The third step is the protein topology prediction, which consists in finding cytosolic, transmembrane, and extracellular regions. To this aim, we used an ameliorated version of Python-TMHMM, which is based on a hidden-Markov model [20]. We implemented the source code available at https://github.com/dansondergaard/tmhmm.py adapting it to work with thousands of FASTA protein sequences. *TMhelices* predicts, for each protein, the number of transmembrane helices domains (TMDn), as well as “Tmhmm seq”, a reduced-alphabet protein sequence containing all different topologies detected, which is crucial for *LoopRazor*. The substitution of HMMTOP [7], the topology predictor used in NERVE 1.0, simplified the installation procedure.

### 2.4. Loop-Razor module

*Loop-Razor* allows the user to retrieve peptides of PVCs, which have TMDn ≥3. It has been introduced because most of the filtration - RV programs, such as NERVE 1.0, discards proteins with TMDn ≥3. Such cut-off has been applied so far to avoid impaired expression of the recombinant protein, a very frequent outcome when dealing with membrane proteins, despite several, recently improved protocols [28,29]. In the pioneering work of Pizza et al. [2], 250 out of the 600 surface proteins of *Neisseria Meningitidis Group B* - endowed with multiple transmembrane domains - were excluded from subsequent characterization steps for unsuccessful cloning. Nevertheless, transmembrane proteins very often turned out to be PVCs because they belong to the surface sub-proteome and hence have more exposed epitopes. To avoid discarding such potentially useful epitopes, as suggested by Olaya-Abril et. al [30], protein fragments that are not embedded in the membrane can be selected for cloning and vaccine testing. To this purpose, when *Loop-Razor* is active, only the outside loops (o-loops) of transmembrane proteins are considered for all proteins with TMDn ≥ 3. With outer membrane proteins (OMPs), the internal loops (i-loops) facing the periplasmic space, are also examined, because they belong to the surface proteome too. In these selected loops, if the longest of a minimum 50 amino acids - continuous sequence (default value, user-modifiable) is detected, it will replace the entire original protein sequence and will then be analysed by the subsequent modules. Considering o-loops, or OMPs-i-loops, is made possible by *TMhelices*, as stated. Reducing the excessive filtration, *Loop-Razor* recovers promising VCs.

### 2.5. Autoimmunity module

This module compares bacterial proteins with human ones to identify similar regions to prevent low immune responses, tolerance issues or autoimmune reactions in vaccine recipients. To achieve this, it retains the two-step structure from the original NERVE:

1. comparison to the human proteome via BLASTp [8]
2. analysis of found shared peptides (SPs) to look for Major Histocompatibility Complexes (MHC)-I II human ligands.

MHC class I epitopes stimulate T-lymphocytes cytotoxic activity while MHC class II epitopes promote helper T-cell response which is essential for antibody production [42]. Presence of such ligands could induce strong autoimmune reactions. The SPs parameters (minimum length, substitution, and mismatch) have been maintained with the same default values and are still modifiable. A new tunable parameter is the e-value, introduced to regulate the number of hits found with BLASTp [8].

The outdated MHCPEP database [31] was replaced with a file containing human MHC ligands retrieved from the IEDB database [32] consisting of 7473 bacterial linear epitopes. In the “Assay” section of the IEDB.org homepage (Figure 3), all types of experiments with positive outcomes have been considered. No filters have been applied to the “Disease” section, instead. To quantify the similarities between microbial and human proteins, we used the same formula adopted in NERVE 1.0 (see *Select module)*.

**Figure 3.**
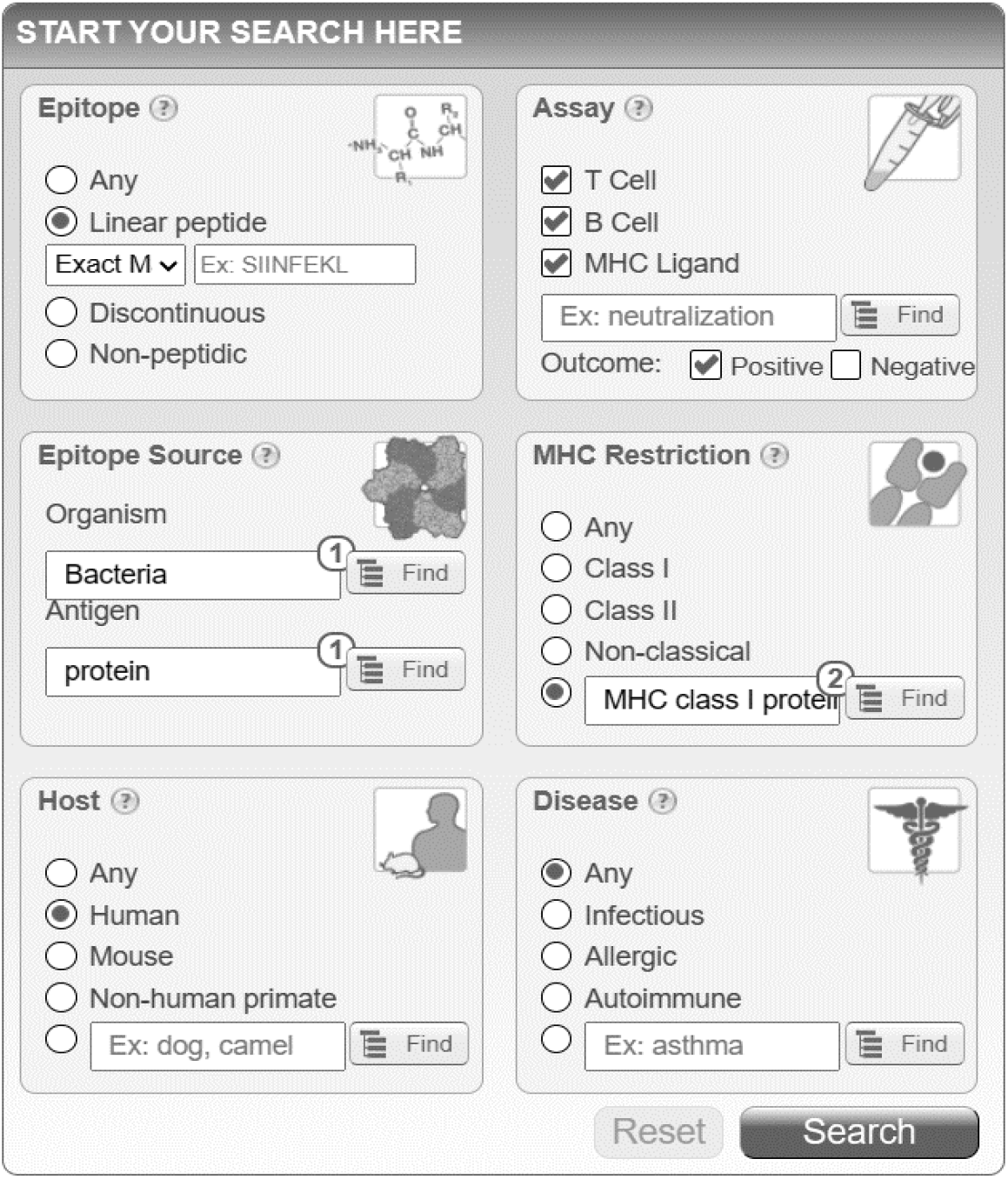
Snapshot taken from the IEDB.org home page. All settings applied to create mhcpep-sapiens are shown. “MHC Restriction” is for MHC class I and II.

### 2.6. Mouse immunity module

This facultative step was added to ease possible vaccine manufacturing - pre-clinical studies. Comparison of input protein queries to the mouse proteome is once again performed by BLASTp. Found SPs are scanned to eventually find matches with mouse MHC ligands. To this purpose, a database of 2060 mouse ligands (mhcpep-mouse) was retrieved from IEDB.org. The applied filters on the IEDB webpage are the same used for *Autoimmunity*, except for the “Host”, which is Mouse in this case). The SPs parameters are also modifiable in this module.

### 2.7. Conservation module

*Conservation* has maintained the same features as in NERVE 1.0. It compares a selected bacterial proteome with another one from the same strain/serogroup using BLASTp. The assumption on which this comparison is based, taken from the original article, is: “*the more a PVC is conserved, the more protective the vaccine produced*” [4]. Ranking of PVCs in descending order of conservation is possible with the formula adopted in *Autoimmunity* and *Mouse immunity* modules.

### 2.8. Virulent module

This newly developed optional component predicts the probability of being a virulence factor (PVR). This overcomes a main limitation of NERVE 1.0, which only considers adhesins and adhesin-like proteins as PVCs [3]. Indeed, even though these two latter categories often represent relevant vaccine targets, other protein classes should be examined as well. An example are invasins and toxins, which are both pathogenic and antigenic proteins, therefore promising targets for vaccine development [11].

It is therefore necessary to consider all protein classes that allow microorganisms to: (i) colonise host tissues, have immunomodulatory and/or suppressive properties, or (ii) deprive the host of essential nutrients which fall under the definition of virulence factors. Since these factors can lack, or have poor, adhesive properties, *Virulent* solves the gaps left by the search of adhesins alone.

To achieve this purpose, we designed a ML model with the same architecture of ESPAAN. Both training and validation datasets were retrieved from the Virulence Factor Database (VFDB) [33] (specifically, the protein core dataset) and from the SwissProt section of UniProt, using keyword search [19]. The following keywords were excluded from the search: *virulent, pathogen, lethal, adhesin, adherence, biofilm, toxin, endotoxin, exotoxin, enterotoxin, invasin, antiphagocytic, motility, flagella, pilus, multidrug, subtilisin, immunoevasion, immunomodulation, lipopolysaccharide, lipoprotein, spore, antibiotic*. After redundancy removal with CD-Hit (25% identity threshold) [25], 1820 virulent factors and 1808 non-virulent factor proteins were left. Then, sequences matching proteins used to tune *Select* have been discarded.

A PCA was applied to perform data reduction, selecting only 400 features. Virulent was trained for 110 epochs and its evaluation metrics are reported in the following Figure 4 and Table 3.

**Table 3.** Evaluation metrics of Virulent calculated on its test set, using a dedicated python script with TensorFlow.

| Metric | Value |
| --- | --- |
| Accuracy | 0.751 |
| Sensitivity (Recall) | 0.724 |
| Precision | 0.768 |
| F1-score | 0.745 |
| False Positive Rate (FPR) | 0.222 |
| False Negative Rate (FNR) | 0.276 |
| True Negative Rate (TNR) | 0.778 |
| False Discovery Rate (FDR) | 0.232 |
| Negative Predictive Value (NPV) | 0.735 |

**Figure 4.**
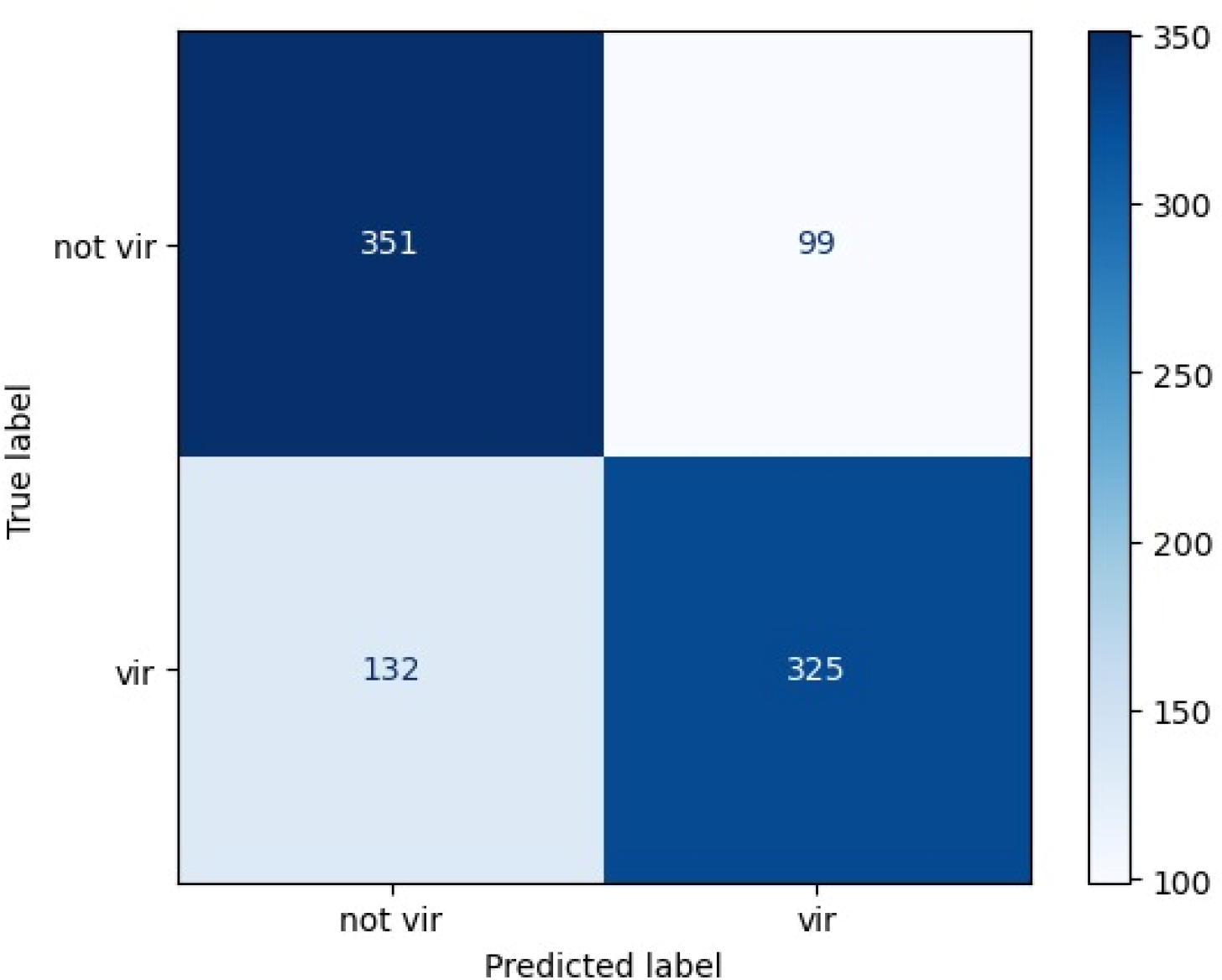
Virulent confusion matrix. Similarly to ESPAAN, here is a 2×2 matrix considering PVR (probability of being a virulence factor) = 0.50 as threshold.

Differently from ESPAAN, Virulent shows reasonable, but not outstanding, performances. Precision and recall have values greater than 0.7. False positive and negative rates are not exactly negligible as the ESPAAN ones, in particular FDR, which is 0.276. Virulent overall accuracy is fairly valid (0.751).

A 3-fold cross validation was performed for Virulent as well (Table 4). This confirmed the quality of the model.

**Table 4.** 3-fold cross validation of Virulent with related obtained metrics.

| Metric | Value | Std deviation |
| --- | --- | --- |
| Mean validation accuracy | 0.767 | 0.019 |
| Mean validation recall | 0.751 | 0.042 |
| Mean validation precision | 0.779 | 0.012 |

For a comprehensive analysis of Virulent tests, see Additional file 1 in *Supplementary Material* and *Availability of data and materials* sections.

### 2.9. Annotation module

This optional module uses the DeepFRI model to infer protein function [34]. The predictor returns Gene Ontology (GO) terms [35] associated with each protein above the confidence threshold of 0.5, in agreement with DeepFRI documentation. The information obtained is not used by *Select* but can be considered to facilitate the manual screening of VCs.

### 2.10. Select module

The PVCs filtration is accomplished by *Select*, following these rules:

- Being not predicted as cytoplasmic;
- Having a PAD value *> padlimit*\*;
- Having a TMDn ≤2 if *Loop-Razor* is off;
- Having sum of SP with host** database)/protein length < 0.15;
- No match with host ligands database;
- PVR *> virlimit\**, if *Virulent* module is active***.

\**padlimit* and *virlimit* values are user-modifiable;

**the host is human and/or mouse if the related module is on;

***If *Virulent* is on, proteins with either PVR *> virlimit* or PAD *> padlimit* are selected.

We decided to make the activation of *Select* optional. So, when it is not active, all protein information collected is viewable.

A composite score ranging from 0 to 1 is associated with each antigen as it is meant to prioritise the best PVCs by combining the normalised scores of each predictor. The configuration of *Select* and its parameters is detailed in paragraph *2*.*12*.

### 2.11. Epitope prediction module

The *Epitope prediction* module identifies epitopes that can potentially bind to MHC-I/II complexes. These molecules, which are codified by the HLAs (Human Leukocyte Antigens) genes, bind and present to T cells the epitopes generated from the intracellular processing of exogenous pathogenic proteins during infections [44]. Given the vast diversity in HLA alleles along with epitopes tridimensional structure and combinations [41, 44], this module considers only linear epitopes and strategically employs the HLA supertypes. This, as defined by Sette and Sidney [36], simplifies the analysis by considering a set of representative HLA alleles to ensure a good coverage of MHC complexes within the world’s population. Indeed, the MHC-HLA polymorphism allows to cluster them into sets of molecules that bind largely overlapping peptide repertoires [44]. Specifically, the selected HLA alleles encompass HLA-B*44:03, 07:02, HLA-A24:02, *03:01, *02:01, 01:01, and HLA-DRB115:01, *13:01, *11:01, *08:01, *07:01, *04:01, *03:01, *01:01.

The Python package epitopepredict [45] was used for epitopes prediction considering the frequency distribution of most common HLA alleles derived from the Allele Frequency Net Database [43]. The user, guided by epitope percentile values, determines the threshold for considering the best-ranked proteins, identified by *Select*. Notably, this component also identifies promiscuous epitopes, which binds to multiple alleles simultaneously. This strategy aims to maximise vaccine effectiveness in the single immunised individual and by increasing population coverage. Moreover, the user is free to personalise predicted epitopes length through specific parameters, including the overlap.

### 2.12. Tuning and benchmarking tests

To tune *Select* and benchmark NERVE 2.0 with state-of-the-art web-RV tools, a dataset containing 615 known bacterial protective antigens (BPAs) was derived from the literature [9,12,37-39]. Proteins were mapped to Uniprot [19] accession codes and sequence redundancy was removed (90% identity threshold) with CD-Hit [25]. Intracellular proteins are not considered as VCs. Therefore, antigens annotated in Uniprot (version 2023-01) as cytosolic or of unknown localization were excluded. Moreover, since PSORTb 3.0 is a NERVE 2.0 workflow - component, BPAs with local identity *above* 25%, as measured using Blast [8], with proteins from the PSORTb 3.0 training dataset were also excluded to prevent data leakage during these procedures. 153 BPAs were obtained, and the dataset was split into 108 antigens for tuning and 45 for testing and benchmarking. Proteomes and organisms of origin associated with each protein were retrieved from Uniprot. Where no proteome was available, antigens were manually added to the reference proteome of the species. To evaluate *Select* performance during the tuning, the fold enrichment was defined as follows:

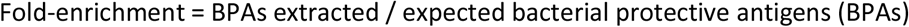

where:

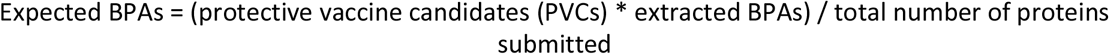

additionally, the recall was calculated as follows:

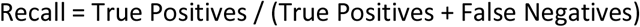

High fold-enrichment values indicate good performances. A hypergeometric test from the python module Scipy (https://scipy.org) was applied to verify statistical significance, as a standard procedure already adopted to test RV programs [9].

It was not possible to obtain a running version of NERVE 1.0 due to its unavailable Perl libraries. NERVE 2.0 was therefore benchmarked against its previous version using the original NERVE 1.0 test dataset, containing 29 BPAs [4]. This dataset was used only for this purpose because it contains several proteins with high similarities with other ones from the PSORTb 3.0 training dataset.

### 2.13. Website building methods

NERVE Web is an open access software with a web interface and distributed computing mechanism on top of NERVE. The server software is written in Node.js and runs on an Express.js server connected to a Redis database and S3 compliant object storage. Docker is used for container virtualisation. The front-end development is written using the Angular framework.

When a processing job is started, an available cluster node starts to run NERVE with input provided. When a job is finished, the user is notified by email and it can view the PVCs filtered with all their analysed features and download the output file.

Created jobs and related IDs are stored locally in the browser, so there’s no need to create an account.

## 3. Results and Discussion

In this section, a focus on the website structure, with the site’s pages descriptions, is provided. A detailed analysis on NERVE benchmarking tests is reported afterwards.

### 3.1 Website interface

To validate our design choices, both the interface and available functions were tested by volunteers not involved in the project (see *Acknowledgements*) and improved based on their feedback. Hereafter we report on individual sections and the functionality of the web application.

#### 3.1.1. Overview and home page

By clicking on: https://nerve-bio.org/, a user enters the NERVE website homepage (Figure 5).

**Figure 5.**
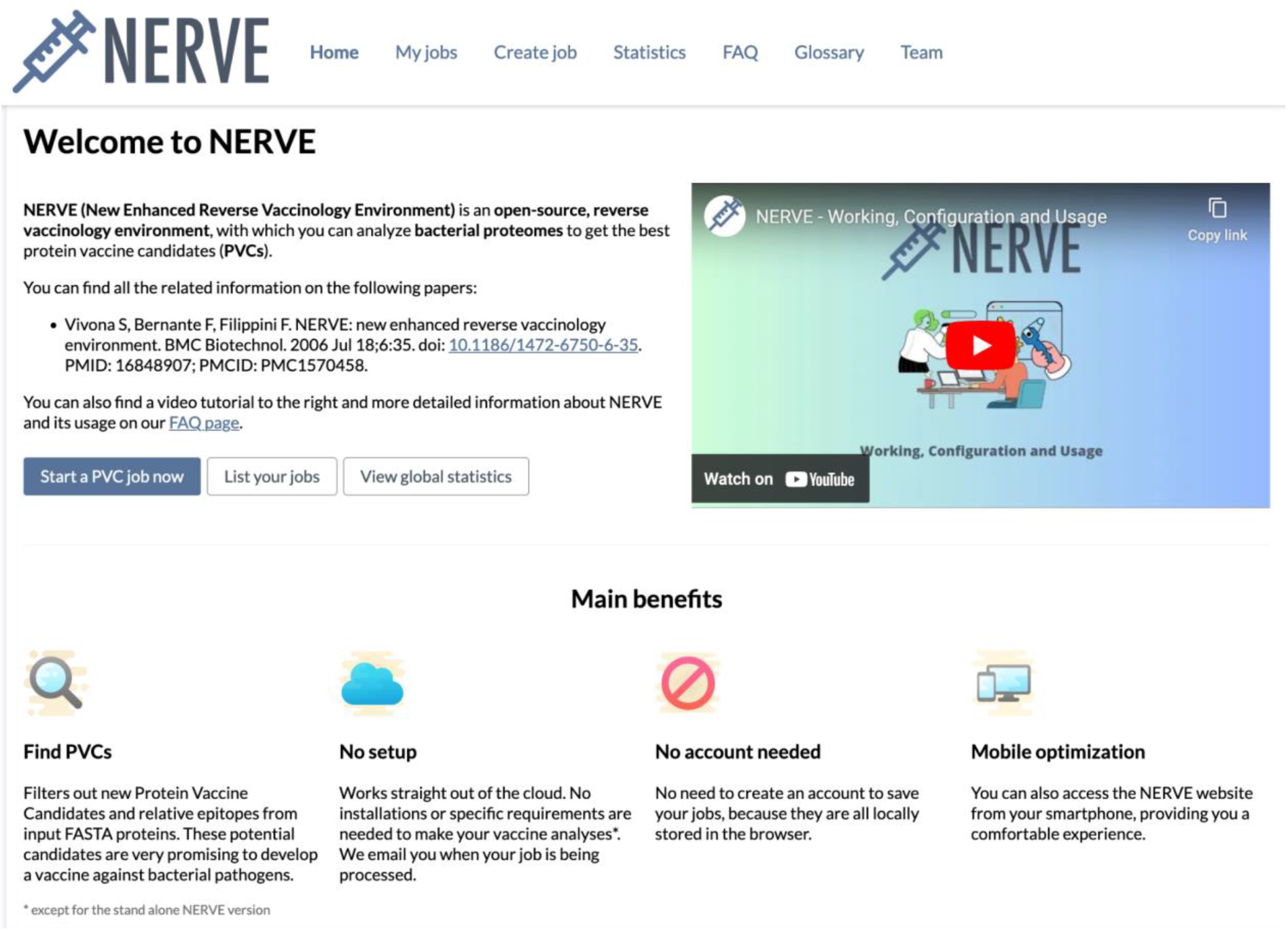
Snapshot from the NERVE 2.0 homepage, with menu header, main benefits, and the dedicated tutorial video.

In the menu header, there are the links to the seven pages of the site: *Home, My jobs, Create job, Statistics, FAQ, Glossary* and *Team*. In *My jobs*, all analyses launched by the user are shown, with the possibility to check their status or to look at the results. Clicking on *Create job*, the user can start a new analysis. The *Statistics* section shows the numbers of user-completed, failed, or delayed jobs. In *FAQ*, NERVE features and working are described along with a tutorial. In *Glossary*, a list of bioinformatics related words, with their definitions, are reported to guide the beginners through their first vaccine analyses. Finally, in *Team*, there are the profiles of people who are contributing to the NERVE project.

#### 3.1.2. Create job

In this section the user uploads its input and sets all the options and parameters for its analysis (Figure 6).

**Figure 6.**
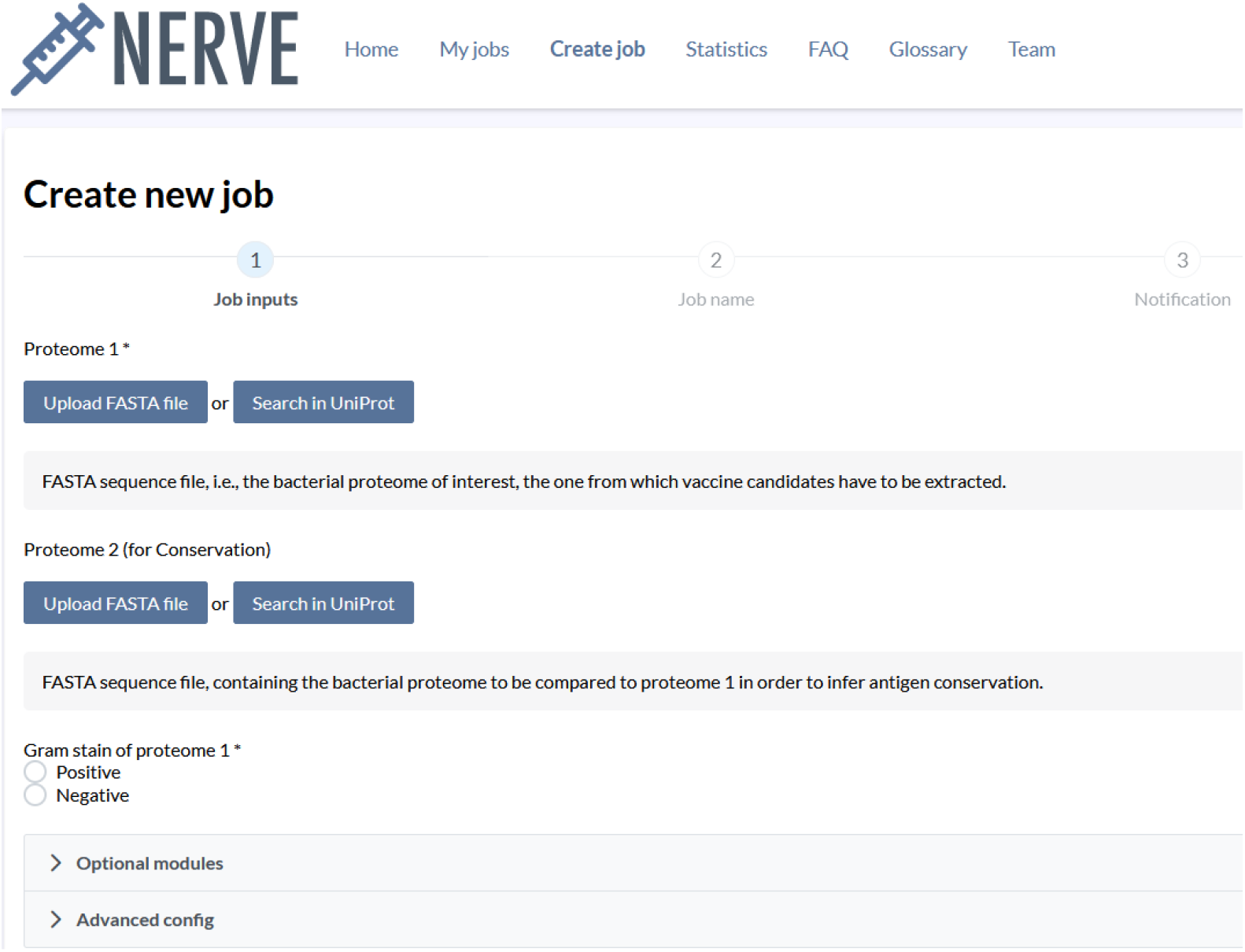
Snapshot from NERVE 2.0 website (Create new job section).

#### 3.1.3. Results visualisation and download

An example of how job results are visualised is shown in Figure 7, 8 and 9.

**Figure 7.**
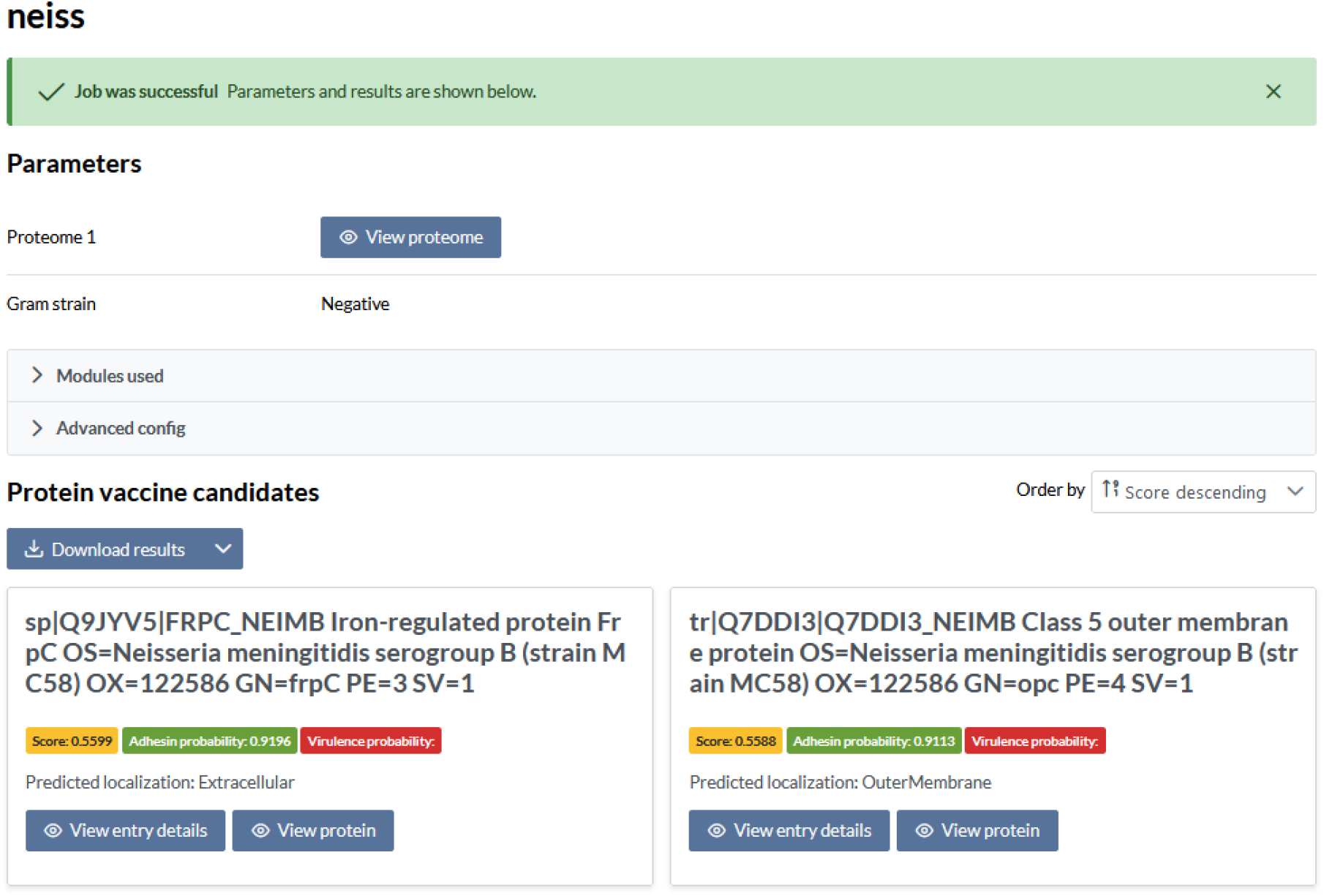
Snapshot from NERVE 2.0 website. Results with PVCs filtered from the proteome of *Neisseria meningitidis* group B (MC58).

**Figure 8.**
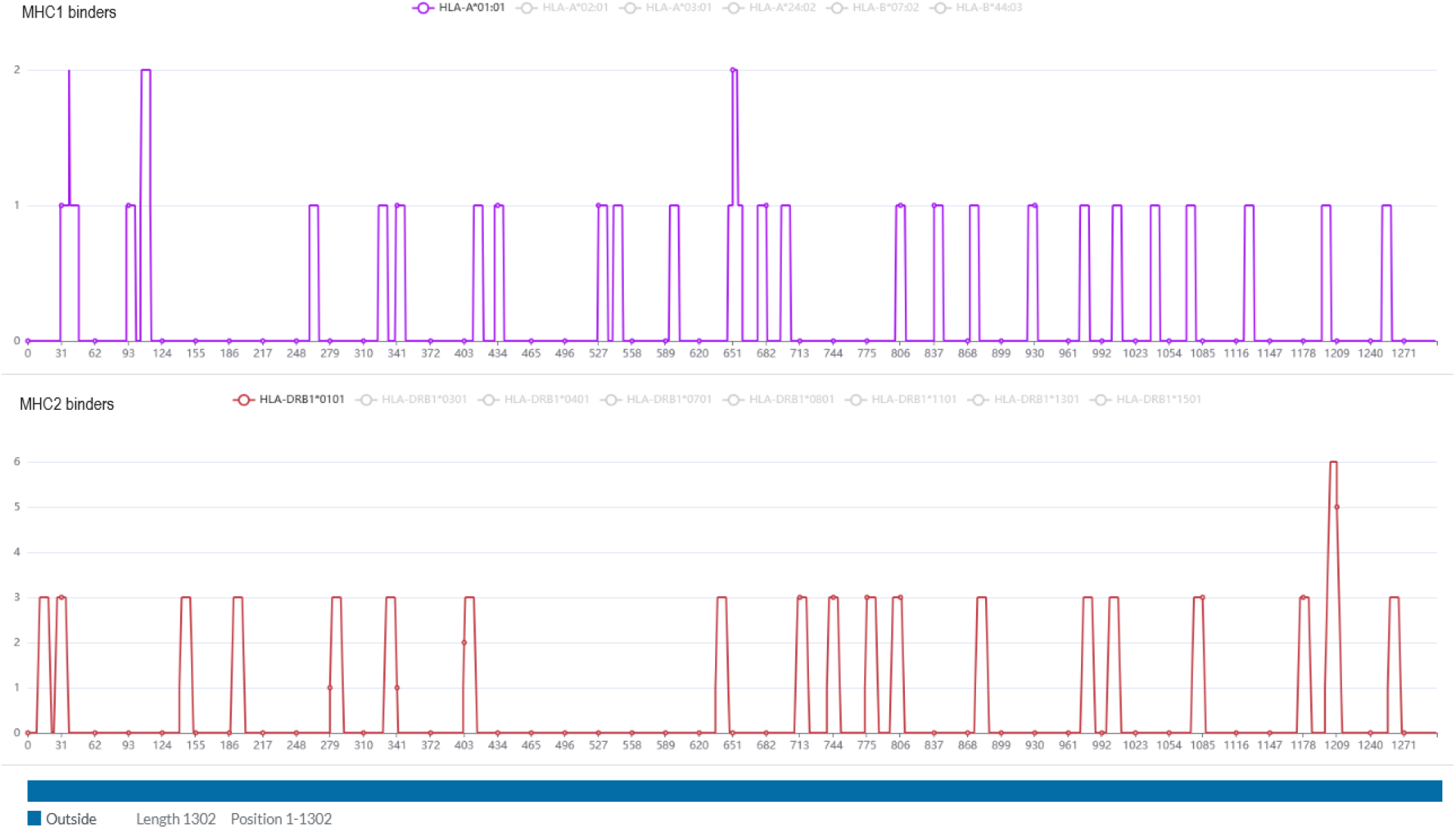
A detail of Epitope prediction results and TMHMM sequence (at the bottom) of UniprotKB: Q9K0K9 extracted from the proteome of *Neisseria meningitidis* group B (MC58).

**Figure 9.**
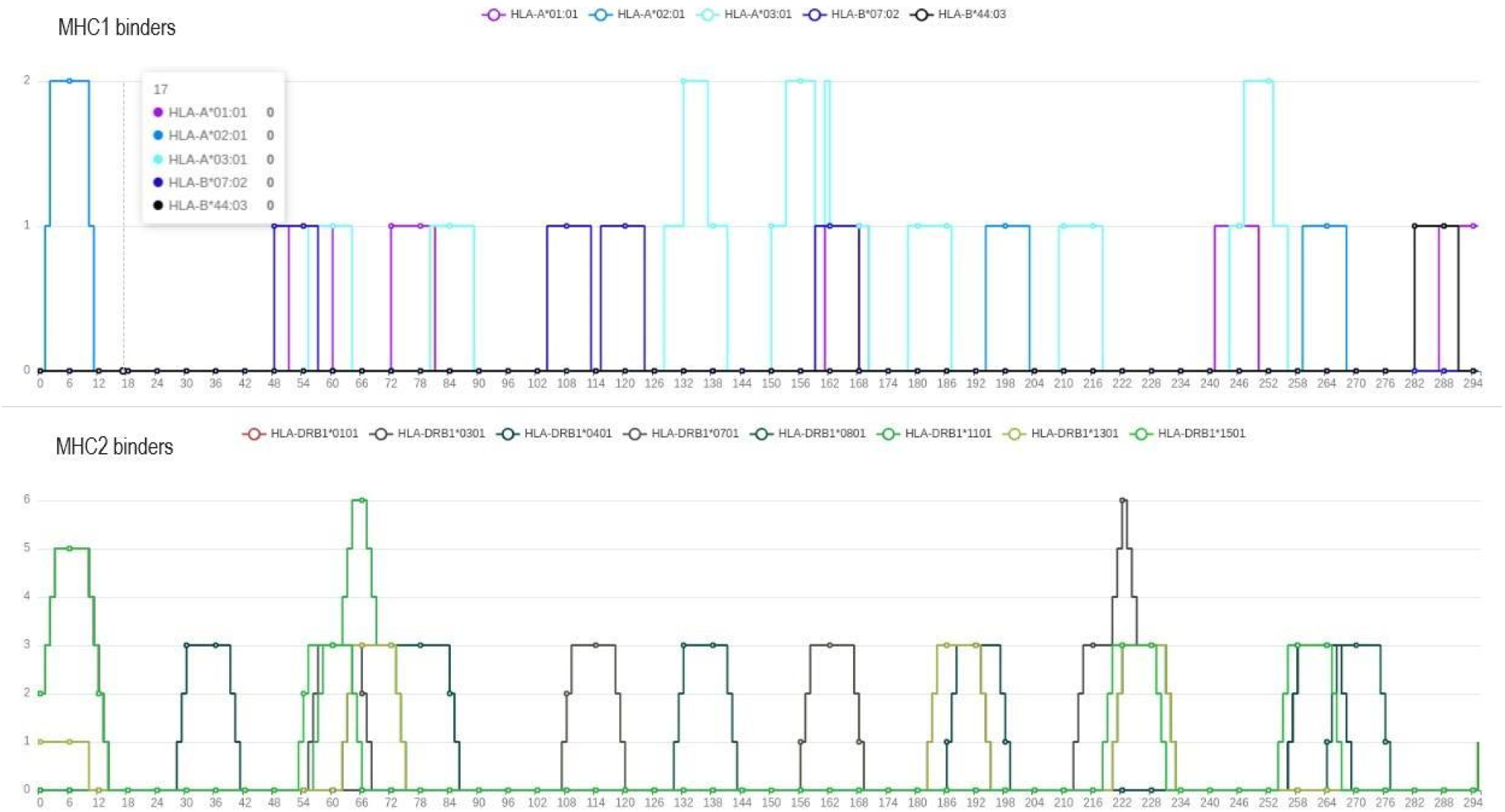
Epitope prediction results for UniprotKB: P17739 from the proteome of *Borrelia burgdorferi (strain ATCC 35210)*. Here are highlighted the MHC-I and II binders for multiple alleles to identify promiscuous linear epitopes.

A summary of all settings and components activated is provided at the top of the webpage. In the *Protein vaccine candidates’* section (Figure 7), the user can see all data collected for each filtered protein by clicking on its related *View entry details*. Here, there is also the Tmhmm seq and the *Epitope prediction* results, with MHC-I and II binders (Figure 8 and 9).

### 3.2. Benchmarking

To find the best *Select* configuration, we tested three different settings on the tuning dataset described in section 2.12: (i) without *Loop-Razor, Mouse immunity* and *Virulent*, (ii) with *Loop-Razor* only, (iii) with *Loop-Razor* + *Mouse immunity* and (iv) with *Loop-Razor* + *Mouse immunity* + *Virulent* combination. For each configuration, the best *padlimit* and *virlimit* thresholds were obtained by performing a 4-fold cross validation procedure resulting in 0.5 for both, which were thus set as default values. The immunogenic and TMDn thresholds were not changed from NERVE 1.0 (see section 2.10).

We evaluated each configuration of *Select* on the test dataset by measuring the fold-enrichment and applying the hypergeometric test.

As shown in Table 5, the best performance was (i) *Loop-Razor, Mouse immunity* and *Virulent* deactivation, (fold enrichment = 16.57). The activation of *Loop-Razor* and both *Loop-Razor* and *Mouse immunity* produced the same results consisting in a fold-enrichment reduction (10.13), while the additional activation of *Virulent* further decreased the fold-enrichment (2.97).

**Table 5.** Tests and benchmark results of NERVE 2.0: highest fold-enrichment values indicate the best performance. 45 known BPAs from 21 proteomes were used and 52,752 proteins derived from the respective proteomes, 36 of which were discarded by NERVE2 after failing quality control analysis. The same dataset was tested on Vaxign2, which failed on performing the computation on 607 proteins, and on Vaxijen. P-values have been calculated with a hypergeometric test.

| Online RV tools | Modules/methods used |  |  | PVCs | Observed BPAs | Recall (%) | Expected BPAs | Fold-enrichment | p-value |
| --- | --- | --- | --- | --- | --- | --- | --- | --- | --- |
|  | razor | mouse | virulent |  |  |  |  |  |  |
| NERVE 2.0 | no | no | no | 796 | 11 | 25.00 | 0.66 | 16.57 | 4.21e-11 |
|  | yes | no | no | 828 | 7 | 15.91 | 0.69 | 10.13 | 5.29e-6 |
|  | yes | yes | no | 828 | 7 | 15.91 | 0.69 | 10.13 | 5.29e-6 |
|  | yes | yes | yes | 4,441 | 11 | 25.00 | 3.70 | 2.97 | 8.34e-4 |
| Vaxign2 | ML-based |  |  | 27,227 | 36 | 81.82 | 22.71 | 1.58 | 3.10e-5 |
|  | filter-based |  |  | 1,059 | 8 | 18.18 | 0.88 | 9.06 | 2.40e-6 |
| Vaxijen | ML-based |  |  | 10,905 | 31 | 70.45 | 9.10 | 3.41 | 1.67e-12 |

Performances were benchmarked against existing web-based RV tools on the same test set: Vaxign2 (https://violinet.org/vaxign2) [12] and Vaxijen (http://www.ddg-pharmfac.net/vaxijen/VaxiJen/VaxiJen.html) [13]. Vaxign2 results were retrieved with a customised Python script for web scraping while Vaxijen results were provided by the authors. We used a web scraping approach for Vaxign2 as its command line version was failing the process for some proteomes with cryptic errors, preventing us from running the computation on 607 proteins. Tests were performed on the remaining set. Vaxign2 was tested using either the score obtained with VaxignML (0.9 threshold), a ML model based on eXtreme Gradient Boosting trained to predict BPAs [40] and, with the filtering method (adhesin probability *>* 0.51, number of transmembrane segments *<*2 and extracellular localization as suggested by the website default parameters). Vaxign-ML showed poor performance, predicting a consistent protein amount as BPAs in the dataset (fold-enrichment: 1.58) while the filtering method performed significantly better (fold-enrichment: 9.06). Vaxijen is also based on a ML method and provides a score associated with each protein.

Proteins with a score *>* 0.4 were considered BPAs according to Vaxijen website. Its performance is better than ML-based Vaxign2 but worse than filter-based Vaxign2 (fold-enrichment: 3.41).

NERVE 2.0 also outperformed its previous version, showing a 38.5-fold enrichment (p-value=1.89e-41) compared to 8.14 of NERVE 1.0 (p-value=8.85e-29) when tested on the NERVE 1.0 test set.

Overall, NERVE 2.0 demonstrated its superior performances compared to both web-based state-of-the-art predictors and to its previous version.

## 4. Conclusions

NERVE 2.0 is now available in a new guise, with a simple and clear web interface to be easily and readily usable. So, user-friendliness represents one of the most relevant features of this significant update. Moreover, new components, with related AI models, and adjustable parameters guarantee respectively an improved and a customisable computational vaccine analysis, meeting the demands of all kinds of users.

NERVE 2.0 showed better performances compared to its predecessor and to other web-based RV programs (Vaxign2 and Vaxijen). Even if the activation of some components may result in lower fold-enrichment values, this is compensated by evidence that these values are still high, while providing the users with new functionalities and minimising the stringency in selection. The user can narrow the extraction of VCs by setting high stringency. Alternatively, it is possible to choose between more candidates that would have been otherwise discarded, as discussed for the Loop-Razor module, which recovers antigens fragments, or for Virulent, which selects further possible VCs when activated.

Together with the web application, the NERVE 2.0 stand-alone version allows the users to perform high-throughput analyses, not being limited to server requests or bad Internet connection. For its installation, it requires: a Linux-operating system, Docker and a few instructions to follow.

Concerning future perspectives, we will further support NERVE with steady improvements and additions. Thus, progressively optimised AI models for protein analyses will be implemented to provide the users with all the necessary tools to refine their vaccine research.

### Availability and requirements

- **Project name**: NERVE 2.0
- **Project home page**: https://nerve-bio.org
- **Stand-alone version**: https://github.com/nerve-bio/NERVE
- **Operating system**: Platform independent. Stand-alone version: Linux operating system.
- **Programming language**: Python
- **Other requirements**: for the stand-alone version requirements see the GitHub page.
- **License**: MIT license
- **Any restrictions to use by non-academics**: see https://github.com/nerve-bio/NERVE?tab=MIT-1-ov-file

## List of abbreviations

RV: Reverse Vaccinology
VC: Vaccine candidate
PVC: Protein vaccine candidate
ML: Machine learning
AI: Artificial Intelligence
ANN: Artificial Neural Network
PAD: Probability of being an adhesin
TMDn: Number of transmembrane helices domains
OMP: Outer membrane protein
BPA: Bacterial protective antigen
MHC: Major Histocompatibility Complex
SP: Shared peptide
PVR: Probability of being a virulence factor
HLA: Human Leukocyte Antigen
FPR: False Positive Rate
FNR: False Negative Rate
TNR: True Negative Rate
FDR: False Discovery Rate
NPV: Negative Predictive Value

## Declarations

### Ethics approval and consent to participate

Not applicable.

### Consent for publication

Not applicable.

### Availability of data and materials

ESPAAN and Virulent GitHub pages: https://github.com/nicolagulmini/spaan

https://github.com/nicolagulmini/virulent_factor_classification

Data about testing their performances:

https://github.com/nerve-bio/NERVE/tree/main/models_data

*Tmhelices* Github page: https://github.com/nicolagulmini/tmhmm.py

NERVE stand-alone version: https://github.com/nerve-bio/NERVE

List of antigens for test and tuning:

https://github.com/nerve-bio/NERVE/blob/main/database/antigens/test_antigens_summary_v2.xlsx

Results of NERVE benchmarking:

https://github.com/nerve-bio/NERVE/tree/main/tuning_and_benchmarks

### Authors’ Contributions

Conceptualization, A.C.,N.G.,F.C.,M.C. and F.F.; methodology, N.G.,F.C.,F.B. and F.P.; software, N.G.,F.C.,F.B.,F.P. and A.C..; validation, A.C.,F.C. and N.G.; formal analysis, A.C.,M.C.,F.C. and F.P.; investigation, A.C., F.C., M.C., F.P.; resources, F.C. and N.G..; data curation, A.C., N.G., F.B. and F.C.; writing— original draft preparation, A.C.,F.C. and N.G.; writing—review and editing, F.F.; supervision, F.F. All authors have read and agreed to the published version of the manuscript.

## Acknowledgements

We thank Alex Bateman, Sandro Vivona, Regina Tavano and Irene Righetto for testing NERVE 2.0, providing us useful feedback.

## Supplementary material

### Additional file 1

NERVE 2.0 supplementary material, with theoretical background of the adopted ML model performance measures.

### Additional figure 1

NERVE 1.0 pipeline, with related caption describing all tools involved and its overall working scheme.

## Additional file 1

### NERVE 2.0: boosting the New Enhanced Reverse Vaccinology Environment via artificial intelligence and a user-friendly web interface

#### Model performance measures

To assess the performances of the two newly-developed neural networks (ESPAAN and Virulent), for each of them, a confusion matrix was created and the most common evaluation metrics were directly inferred from it. Additionally, a k-fold cross validation was used to evaluate how the models work with new unseen data. Therefore, in this section, a brief theoretical background of these mentioned model performance measures is reported.

##### 1. Confusion Matrix

A confusion matrix can be defined as an organized way to summarize the performance of a ML (machine learning) model [1], showing which of its predictions are correct and which incorrect for all classes. In this specific case, each model classifies its input in two distinct classes.

In fact, both ESPAAN and Virulent have to solve a binary classification problem. The first model classifies input-analyzed proteins into adhesins or not-adhesins; Virulent, instead, predicts if a protein is a virulence factor or not.

Since dealing with two binary classifiers, we created two bidimensional confusion matrices. A common “2×2” confusion matrix, reported as an example, is shown in Figure 1.

**Figure 1:**
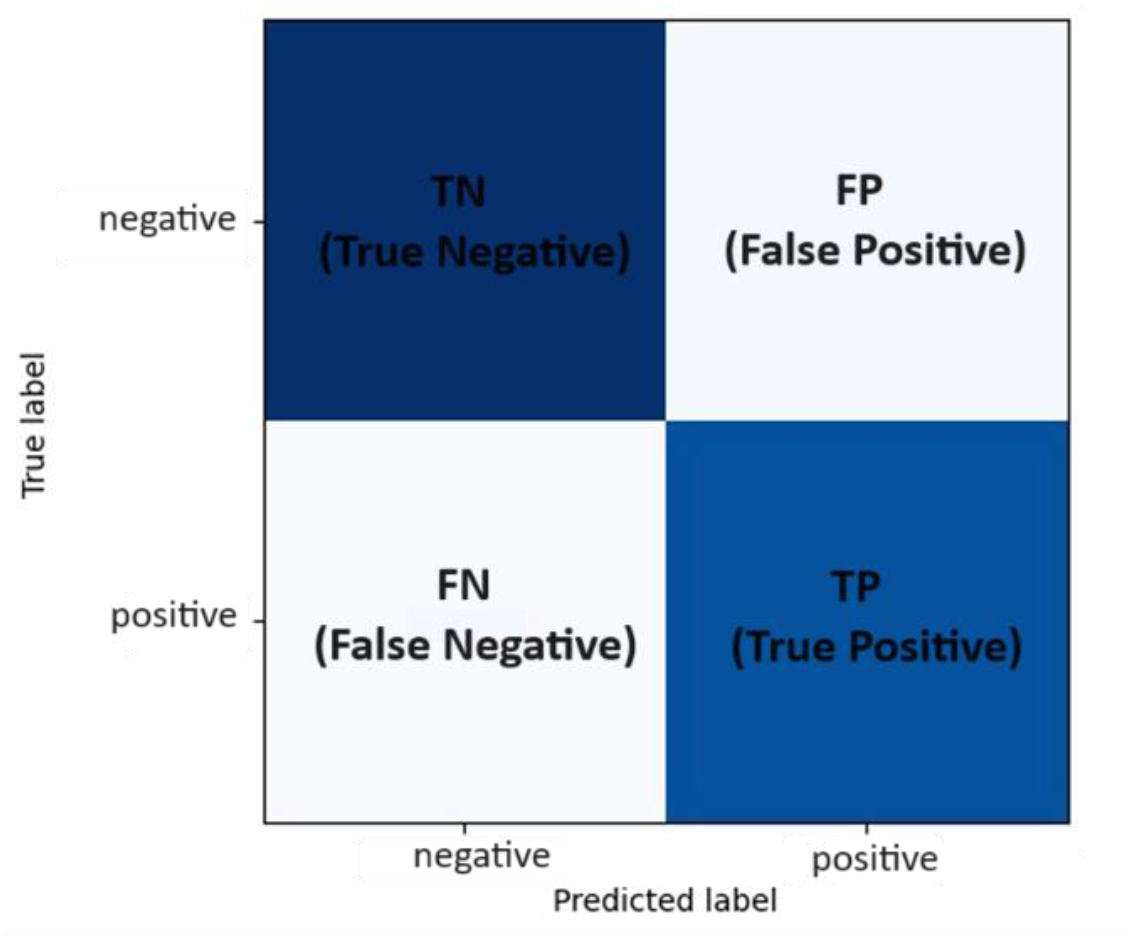
Working scheme of k fold cross validation with k=3

As evidenced, there are four different values (one for each matrix cell), which are useful to understand if the tested model is making predictions which deviate from real values, or not, and how many times this happens. It is necessary to underline that these real values come from the known datasets we used to train our binary classifiers (supervised machine learning).

The values mentioned in Fig. 1 are:

- TN (True Negative), which is the number of real negative samples that the tested model classifies as negative.
- TP (True Positive), which represents the number of real positive samples correctly classified.
- FP (False Positive), which is the number of real negative samples, classified as positive.
- FN (False Negative), which is the number of real positive samples, classified as negative [1,2].

So, in the case of ESPAAN, TN is the number of known non-adhesins which are correctly identified by ESPAAN as non-adhesins. Instead, FP in Virulent represents the number of known non-virulent proteins which are classified as virulent.

From these values, it is possible to compute the model evaluation metrics.

##### 2. Evaluation metrics

The evaluation metrics are crucial to assess the effectiveness of ML models, being quantitative measures of their performance [2]. All their values are expressed with a number between 0 and 1. We used nine different metrics here explained.

One of the most used metrics for classification is accuracy, which is calculated with the following formula:

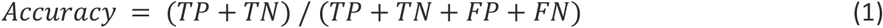

It’s a fundamental metric representing the number of all correct predictions out of all predictions. So it shows the overall correctness of the ML model [2].

Another important metric is the precision, also called Positive Predictive Value (PPV). It shows the correctness of all the positive predictions, so it is the portion of TP among all model positive predictions [2]. And it is

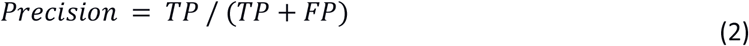

defined as:

Recall instead, known also as sensitivity or True Positive Rate (TPR), is the model’s ability to find all the positive samples in a dataset and it is expressed as the ratio between TP and all actual positives (TP+FN) [2].

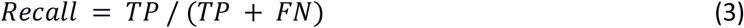

The harmonic mean between precision and recall values, obtained by combining them, is known as F1-score [2].

A low F1-score often indicates an imbalanced performance of the ML model. On the other hand, a F1-score closer to 1 indicates that the model, with high recall and high precision, is really good at recognizing both positive and negative samples. This score can be computed adopting this formula:

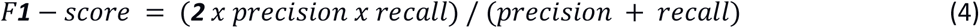

Another considered evaluation metric is Specificity or True Negative Rate (TNR). As shown by the following formula, it is the portion of TN that was correctly identified in the chosen dataset.

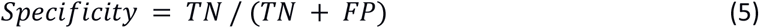

This measure describes how well a ML model can identify TN [2].

Instead, Negative Predictive Value (NPV) is the portion of TN among all model negative predictions.

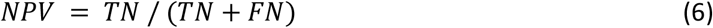

This value can also be seen as the opposite of PPV, or precision. Indeed, precision measures the TP fraction over all positive predictions, instead NPV is focused on TN detection from all negative predictions [3].

The last three reported measures can be derived from some of the just analyzed metrics. Differently from the latters, when evaluating a good-performing model, low values of these rates are expected.

The False Positive Rate (FPR) is calculated as the ratio between FP and all actual negatives.

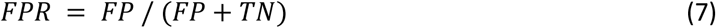

It can be also calculated with this formula: *FPR* = *1* −*TNR* (or specificity) (8) [3].

The False Negative Rate (FNR), also called miss-rate, is expressed as the ratio between FN and all actual positives.

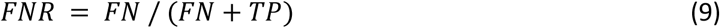

Another usable formula for FNR is: *FNR* = *1* − *TPR* (or recall) (10) [3].

Lastly, False Discovery Rate (FDR), is the fraction of FP over all positive predictions and it can be obtained as follows:

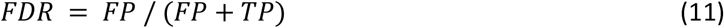

FDR is also complementary to PPV, or precision. Indeed, it can be expressed as: *FDR* = *1* − *PPV* (12) [3].

Evaluation metrics values for ESPAAN and Virulent are listed and commented respectively on 2.2 and 2.8 paragraphs in the article.

##### 3. k-fold cross validation

K-fold cross validation is a specific type of cross validation (CV) technique, which is widely used to test ML models performance. In particular, it can be really helpful in avoiding overfitting, an undesired situation which occurs when a ML model has great accuracy with training data but not with test data. Then, k-fold CV is useful to estimate the model generalization, and so its ability to do appropriate predictions adapting to new input data yet to be tested [4].

As shown in Figure 2, the dataset is divided into *k* approximately equal parts or “folds”.

The tested model is trained *k* times using a different fold as the validation set and the remaining *k-1* folds as the training set. This way, every data point in the dataset is used once for validation and *k-1* times for training. The value of *k* can vary, with common values being 2, 3, 5, 10, or even 20. The value is empirically chosen, and ideally, the best value is the one that minimizes both the bias and the variance of the model.

In our approach, we aimed to keep a reduced model complexity by diminishing the number of parameters. Consequently, we expected a high bias and low variance in our results. Achieving strong performance in both training and validation phases—using our established Keras validation framework for training the current models—we opted for k = 3 to ensure our models ability to generalize. This choice implies that each iteration’s training set closely reflects the original, in cardinality. Increasing *k* would result in smaller training sets per iteration, potentially compromising the models learning process [6][7].

**Figure 2a:**
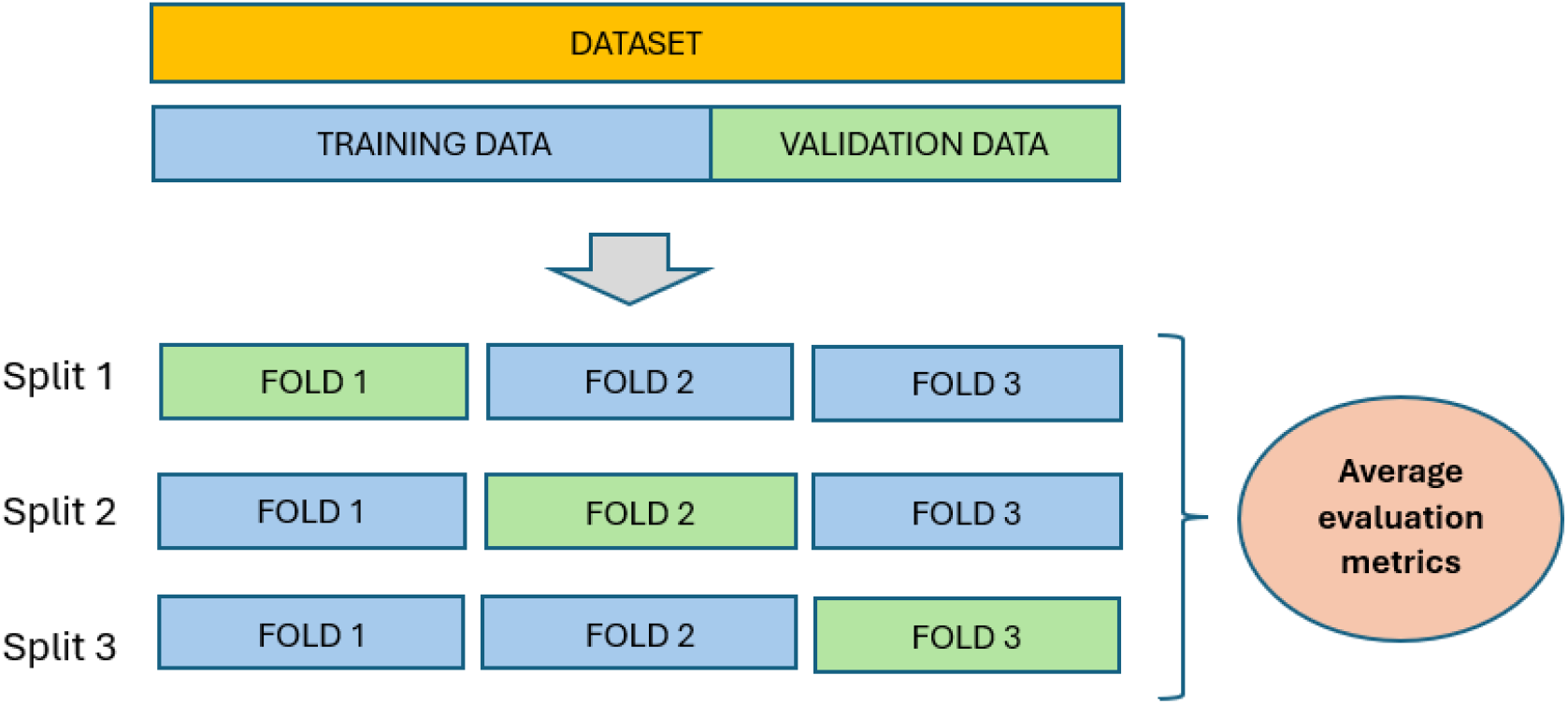
Example of a 2×2 confusion matrix

For each split, the principal model performance measures are computed and the score of the entire procedure is determined by averaging the resulting values [4].

The relative standard deviations have to be computed, as well as the mean values.

Standard deviation (*σ*) is a fundamental statistical tool that measures the dispersion or the variability of a dataset, intended here as a set of measures, around the mean value. It is also useful to compare the variability of datasets having different units of measurement [5]. A smaller value indicates less variability, while a larger one indicates greater variability. In our case, we used the standard deviation to quantify the error/uncertainty connected to model measures, specifically the CV mean evaluation metrics.

It is defined as the square root of the variance (σ^2^), which is calculated using the following formula:

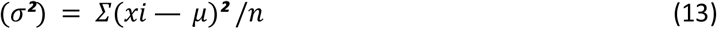

Where *Σ* represents summation, *xi* is each data point, *μ* is the mean of the dataset and *n* is the dataset size (number of data points) [5].

So, the standard deviation is expressed as:

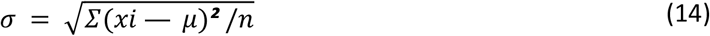

Even if the variance is a dispersion data measure as the standard deviation, it is not generally expressed in the same units as the dataset and doesn’t have smaller values than the second one. All this makes the standard deviation an easier interpretable measure of data dispersion [5].

All k-fold cross-validation mean values and the related standard deviations are reported and discussed in the and 2.8 paragraphs of the article.

---

**Additional figure 1.**
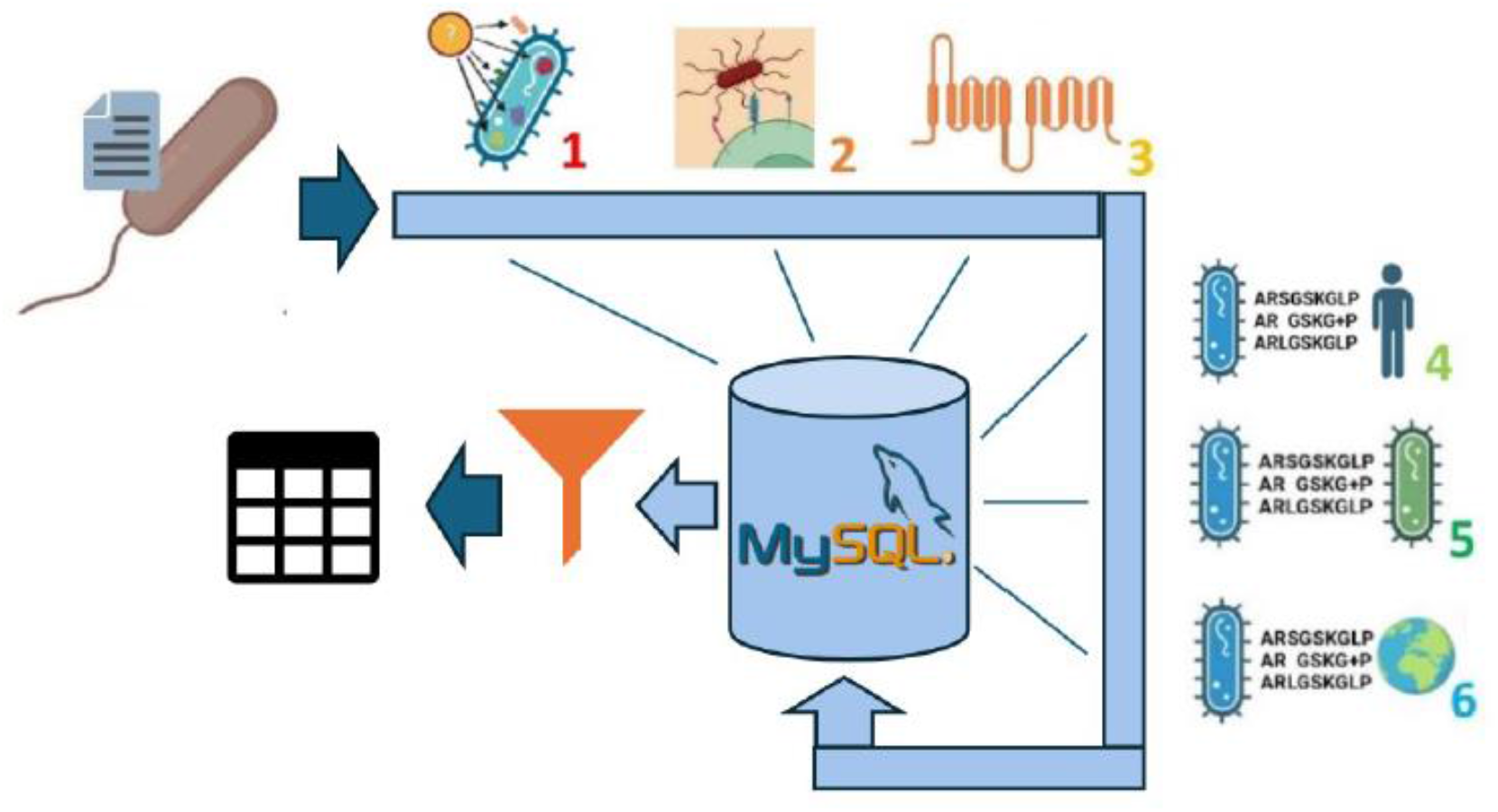
Original NERVE pipeline. Protein sequences from an input proteome are analysed through six steps, including predictions of: (1) subcellular sorting by PSORTb [5], (2) adhesin probability by SPAAN [6], and (3) membrane topology by HMMTOP [7]. Moreover, (4-5-6) sequence comparison applications based on BLASTp [8] are used to identify conserved epitopes and determine inter-strain conservation. Then, the collected information is saved in a MySQL table and finally filtered by the last module, named select. Extracted PVCs are listed in an HTML table. Created with BioRender.com.

## Footnotes

1 https://github.com/kitamura-felipe/adhesin_finder

